# Adverse effects of hunting with hounds on participating animals and human bystanders

**DOI:** 10.1101/2022.08.16.504031

**Authors:** Adrian Treves, Laura Menefee

## Abstract

Hunting mammals with hounds is little studied. We present two datasets consisting of quantitative and qualitative data from self-selected respondents. The first came from hound handlers’ reports of hound injuries with post hoc verifications by government agents. The second came from by-standers reporting eyewitness encounters with hounds or handlers. Self-selected samples cannot be used to extrapolate rates in space or time but do provide nuances of human-animal and human-human interactions. From In the state of Wisconsin, USA, we describe government data on 176 hounds reported to have suffered injury during encounters with wolves. The government did not collect data on wolves or other non-target animals that may have been injured during these encounters. We investigate two wolf-centered hypotheses for wolf-hound interactions, find little support for either, and propose new hound-centered hypotheses. We also describe 105 human bystanders’ reports of experiences with hounds, handlers, and law enforcement agents.

## Introduction

In the face of biodiversity crises and concerns raised about animal ethics partly caused by climate change and partly by human-induced mortality, some societies are re-examining many human uses of animals that once seemed unobjectionable. For example, uses of poison, off-road vehicles, fire suppression, etc. have undergone scrutiny for their societal benefit-cost estimates and their effects on domestic animals, wildlife, and ecosystem health. Among the activities facing renewed scrutiny is hunting with free-running hounds loosed far from their owners. In our search for work on hunting with dogs or hounds in Google Scholar, the search phrase ‘hunt with (hound or dog)’’ yielded 38,300 results declining by half when ‘-bird’ was added to the search string to exclude bird-hunting with dogs. By contrast, ’hunt -bird ‘yielded 3.16 million results. Therefore, hunting with hounds seems under-studied. The gap in information is not only found in Google Scholar but also in legal proceedings judging from one court case in Wisconsin (Brown v Kemp 2021). Namely, the practice of hound-hunting is protected from outside scrutiny such as video-recording or observing hound handlers in public places (Brown v Kemp 2021). Yet, the practice of hunting mammals with hounds has been recorded since 8,000 years at least (Guagnin et al. 2018), praised by President Theodore Roosevelt in 1902 (Roosevelt 1902), and is legal in numerous countries and several U.S. states (Heberlein 2000; Hristienko & McDonald 2007).

Loosing mammal-hunting hounds to pursue prey, some as large as brown (grizzly) bears, *Ursus arctos*, may have harmful effects on people in their path, on the hounds themselves, and on target and non-target wildlife they encounter or pursue (Grignolio et al. 2011; Gompper 2013). Therefore, we present data on reports by self-selected hound-handlers alleging harm to their hounds, following interactions with wolves. We also present self-reported perceptions of human by-standers who experienced the behaviour of hounds during training or hunting and self-reported interactions with handlers or law enforcement agents the bystanders called in after an encounter. Although it is routine in research on carnivore predation on domestic animals to report only adverse encounters, e.g., (Treves et al. 2002), such presentation is clearly incomplete. For an under-studied phenomenon such as hunting mammals with hounds, both adverse and beneficial effects might be studied. The scope of this report is limited to the qualitative and quantitative data from plaints. Therefore, our scope are adverse effects only as with carnivore predation on domestic animals.

### Non-target animals and their interactions with hounds

When pets kill wildlife, biodiversity may diminish or ecosystem health may deteriorate (Grignolio et al. 2011; Gompper 2013). Dogs are potential predators of domestic animals and wild ones (Bowers 1953; Ciucci & Boitani 1998). behavioural changes in wildlife are often reported. For example, American black bears *U. americanus* were reported to avoid encounters with hounds and in so doing approach people and major roads more frequently (Stillfried et al. 2015). Some animals will stand their ground when hounds encounter them (Treves et al. 2002; Wydeven et al. 2004; Backeryd 2007). Hounds are often used for pursuit of mammals larger than individual hounds. When hounds encounter animals that can defend themselves effectively, the hounds may be injured. Larger size and greater competitive ability of the wild animals may alter the risk posed by hounds and vice versa. Researchers have examined aggressive encounters between wolves and dogs in many regions (Fritts & Paul 1989; Backeryd 2007; Kaartinen et al. 2009). The States of Wisconsin and Michigan, USA, have a relatively longer history of such research. Spatial patterns of wolf *Canis lupus* attacks on hounds are somewhat predictable (Treves et al. 2002; Wydeven et al. 2004; Bump et al. 2013; Olson et al. 2014). The risk of an attack appeared to be higher in areas with more public land, larger wolf packs, closer to a wolf pack, and when baits were left out longer. Building on these prior studies, we investigated two hypotheses about wolf-hound interactions (WHI). First, the predation hypothesis predicts WHI occur when wolves attack hounds for food, and second, the territoriality hypothesis predicts that WHI occur when wolves defend territory or pups (Treves et al. 2002; Wydeven et al. 2004). To do so, we describe here the self-reports by owners on the characteristics of both canids involved, and self-reported correlates of the outcomes of WHI. At the time of our study, it was illegal for hounds to attack wild animals. Nevertheless, such attacks might have occurred. We do not have evidence of which animal in a WHI initiated aggression or escalated it to the point of injury or death. We only present data on the outcomes for hounds because outcomes for wolves were not documented. Therefore, we cannot rule out the possibility that wolves responded defensively to hound attacks. The evidence for wolf attacks on hounds came from handlers seeking compensation or other forms of redress (Treves et al. 2002; Ruid et al. 2009; Treves et al. 2009). In a prior study, a number of wolf deaths caused by other canids were invariably attributed to other wolves (Treves et al. 2017b). Yet, veterinary pathologists might not be able to reliably distinguish large dogs, such as hunting hounds, from wolves by simple scrutiny of bite marks without DNA analysis (Plumer et al. 2018). Therefore, our sample is necessarily biased toward handler concerns and outcomes for hounds.

### Hounds and humans

There are no published data on hound-handler’ encounters between by-standers and either handlers or their free-running hounds. There are prior studies of hound handlers’ attitudes though (Supplementary Material 1). In brief, as a group, hound-handlers have the lowest tolerance for wild wolves in any group measured thus far. Yet, interactions between hounds and bystanders or bystanders and hound handlers are not yet documented to our knowledge. The Sierra Club Wisconsin Chapter (SCWC) initiated such a study after a request for information from government agencies and law enforcement was returned without data. Therefore, SCWC members led an effort at gathering information more broadly and systematically.. As part of a National Sierra Club initiative, the SCWC subcommittee solicited first-hand reports members had received of by-stander experience. Because such first-hand reports are likely to be remarkable, most reports were adverse. As with the hound handlers’ self-selected reports, the SCWC survey respondents were self-selected.

Self-selection bias tends to focus on extreme events not average ones. Therefore, spatiotemporal patterns or estimates of frequency in absolute terms would be misleading. We avoid such inferences. However, self-selected reports can describe the nature of interactions and nuances that third parties may not be able to document and also events during interactions that may be invisible to third parties post hoc. Therefore, we offer the reports as a starting point for understanding and managing interactions between hounds, wildlife, bystanders, and handlers.

## Methods

### Wolf-hound interactions (WHI) behaviour, grouping, and phenotype

During our study period, hounds were legally used to hunt many mammals, including smaller carnivores and black bears. Bear hunting in Wisconsin occurred legally from the last week of August into October, and hound training was legal in July and August during the study period (Treves et al. 2009; Treves et al. 2010). Hound hunters accounted for approximately 40% of the annual take by bear hunters (Treves et al. 2010; Olson et al. 2014). Typically hounds were loosed from vehicles and allowed to run far from owners, without direct control (Wydeven et al. 2004). Hounds were often fitted with global positioning systems (GPS) or VHF radio-collars, allowing the owner to follow remotely the movements of hounds and determine when and where a bear had been treed or another animal encountered. Hunters used groups of up to 6 hounds to track and trail prey during training or hunting (Wydeven et al. 2004).

We examined WHI from 7 August 1999 through 19 January 2012. Case files documented 145 killed and 31 injured hounds deemed by state or federal verifiers as “confirmed” (85%) or “probable” (15%) WHI, see methods in (Treves et al. 2002). WHI case files included written reports and sometimes detailed forms documenting field investigations, including necropsy data, photos, veterinary reports, and anecdotal reports from handlers. During the early years of record-keeping, documentation and reporting of depredations lacked uniformity; thus, some portions of the data were missing, resulting in lower sample sizes for various analyses. Although methods of field verification have not been described scientifically, it appears that one or more government agents would receive a complaint and attempt to visit the site of WHI to examine physical evidence. It appears in some unquantified proportion of cases, there was no evidence left but for injuries on hounds. In some of these cases, the injures on hounds were attributed by private veterinarians paid by the owners (Treves et al. 2002). WDNR provided compensation for domestic animals injured or killed by wolves, including hounds (Treves et al. 2002; Ruid et al. 2009; Treves et al. 2009). United States Department of Agriculture agents assumed responsibility for verifying WHI in 1990, and conducted most of the investigations used in our analysis of WHI (Willging & Wydeven 1997). Owners with confirmed losses in Wisconsin were eligible to receive up to $2,500 per hound based on the estimated value of the hound. It has been a well publicized program. Since the compensation program began in 1985, nearly $350,000 dollars were paid to hunters to compensate for hounds injured or killed by wolves. Between 1985 and 2006, payments for hunting hounds comprised 37% of all compensation (Treves et al. 2002; Naughton-Treves et al. 2003; Ruid et al. 2009; Treves et al. 2009). The current working hypothesis is that most WHI incidents were reported because of a compensation program characterized as more generous than other jurisdictions (Naughton-Treves et al. 2003; Treves et al. 2009). Corroborating, Bump et al. (Bump et al. 2013) suggest fewer WHI are reported in Michigan’s wolf range because hound owners receive no compensation.

In WHI files, 91% occurred while pursuing bears, bobcats, or coyote, which involves different breeds and use of hounds than for other quarry, such as waterfowl, upland birds, or rabbits. Hounds that hunt large prey such as bears or coyotes are typically breeds of large size and build, frequently Plott (18-27 kg), or larger hound breeds (Bluetick, Redbone, or Walker coon-hounds (18-36 kg). Occasionally, WHI files did not specify the type of prey being pursued. In these cases, if the breed of hound was one of the five listed above, we assumed the WHI occurred while pursuing the above three wildlife species. In total, we report on 176 case files. We quantified the frequency of WHI among breeds of hounds. If the hound was reported as a mix of multiple breeds, we used the first breed listed. We pooled breeds in an “other” category when a single breed had too few WHI to meet the assumptions of the *chi*-squared test. No data are available on breed frequencies or preference by hunters in Wisconsin, with which we could estimate relative risk by breed.

Our analysis of the body site bitten was limited because a number of hound carcasses were partially or wholly consumed before retrieval by an owner arriving late at the scene. We pooled head, neck, and throat into one category and all other sites in another category, to test if outcomes of WHI differed by bite site. We compared hound group size, number of hounds involved in the WHI, and wolf pack size using (1) number of wolves seen and reported by hunters (observed), and (2) WDNR-reported wolf pack sizes from the winter preceding the WHI (censused). Because wolf packs exhibit fission-fusion sociality and packs disaggregate often in the summer (Mech 1970), when many WHI occurred. Wolf packs in Wisconsin average 4 adults in late winter (range 2-12), usually composed of offspring of the two (usually) unrelated breeders. By summer, pups are exiting the dens but range near them or near other pack rendezvous sites until the pack regularly fuses and travels long distance in mid- to late Fall. During this time, most wolf pack members return periodically to rendezvous or den sites to assist with pup-rearing, and consequently have higher food demands, perhaps requiring wolves to forage more frequently (Ruprecht et al. 2012). From birth until the end of August, wolf pups experience the highest growth rates, with September representing a critical month for weight gain (Kreeger 2003). In some cases, wolf pups have been observed gaining as much as 3.6 pounds per week (Van Ballenberghe & Mech 1975). Pup growth, critical to survival, is limited by food quality and availability. By late August, growth begins to taper (Pulliainen 1965), as does rendezvous site use (Van Ballenberghe & Mech 1975; Kreeger 2003; Ruprecht et al. 2012). The seasonality of wolf life-history on an annual cycle may elevate the risk of WHI in late summer and autumn because of all pack members’ and especially pups’ caloric needs. Finally, we analyzed the temporal occurrence of WHI as it relates to public hunting seasons, and we compared the frequency of WHI during the hound-training period (July-August) to that during the bear-hunting season (approximately September-October, Treves et al. 2010).

### Handler behaviour and husbandry

The state wildlife agency implemented several methods for mitigating or preventing WHI, including compensation for handlers’ self-reported losses, encouraging the use of warning devices on collars to deter wolves, statewide communication to hunters on recent hound injuries and their locations, and designation of Wolf Caution Areas (WCAs). The state communicated the location of higher-risk WCAs online, posted in the field, and in other ways to handlers (Wydeven et al. 2004; Olson et al. 2014). Within WCAs, the WDNR recommended that bear hunters release hounds >2 miles from known rendezvous sites. Prior work documented handlers’ willingness to risk hounds in posted WCAs, even within the same season and even within hours of previous WHI or WCA posting (Wydeven et al. 2004; Olson et al. 2014). Compensation records also document multiple payments to the same owner or handler within a single season (Treves et al. 2009). These data suggest not all hunters heeded the state’s warnings. Handlers may be able to prevent WHI by using protective vests or stronger collars (Backeryd 2007; Khorozyan et al. 2020), keeping hounds leashed until the targeted game species is located, or bringing first aid kits on the hunt, although the possible effect of these interventions has not been studied in Wisconsin.

### Survey of by-standers

The Sierra Club Wildlife Committee (SCWC) subcommittee on Protecting Native Forests and Wildlife collected information from by-standers experiences with hounds loosed during hunting or training. The SCWC posted the survey on the Sierra Club Wisconsin Chapter website 2015-2021, and administered the survey. The survey appeared at https://docs.google.com/forms/d/e/1FAIpQLSd6LcXrzUh-aP061eyQ1hJQTS_8QKulY0IQxCz2WhWVCrNQDw/viewform?c=0&w=1 accessed 1 August 2023. SCWC members also printed hard copies of the instrument and distributed these at wolf and wildlife related meetings and conferences in Wisconsin in 2015 and 2016. SCWC also invited any members of the public who described adverse hounding encounters to fill the online report. Therefore, respondents were self-selected with attendant biases discussed below.

About 80% of respondents used the online form to report anonymously, and 20% sent their responses directly to SCWC via mail, phone, email, or in person while being assured of anonymity. SCWC administrators stripped identifying information from the data and shared the data with LM for publication. We analyzed anonymized data stripped of identifying information. University of Wisconsin-Madison Human Subjects Protection Institutional Review Board states that anonymized third-party data is exempt from review. The 25-question survey (Supplementary Material 2) is organized in four sections: Observations; Trespass; Property Damage, Personal Injury or Threats; and Interactions with Law Enforcement, totaling 22 yes/no questions and 4 items that allowed unstructured responses to elaborate on answers. Respondents could identify county in which the interaction occurred. Respondents could report how many hounds they saw during each interaction. We used the average of the latter data to cross-check with average hound party size from WHI records. When two respondents mentioned the same interaction by date and county, but different numbers of hounds, LM averaged and rounded up for the number of hounds. LM screened the sample to eliminate responses that did not report an interaction but only expressed an opinion about the practice of hunting with hounds (Supplementary Material 2). After the screening, the sample presented here appears to come from independent incidents although we had no way to verify location or date.

We evaluated two prior hypotheses for WHI: territoriality and predation. In brief, territoriality might lead wolves or hounds to interact aggressively, which might be expected to affect seasonal rates of WHI because either canid could have more to protect in certain seasons. For example, wolf packs protect pups while young in summer or early fall. Hounds have regular ranging areas and only part of the year access to those areas. Regarding the predation hypothesis, periods of higher caloric need and lower food availability for wolves might lead to higher rates of WHI in late summer or early fall (caloric needs) or winter (low availability). The state of the carcass might reveal if predation or scavenging were primary or secondary motivations for WHI.

### Statistical analyses

We performed statistical analyses (SAS Institute 2013) using Student’s paired *t* tests to compare the differences in average estimated ages of hounds and numbers of wolves during the attack, in relation to the outcome of the WHI (i.e., killed or injured) after evaluating if variances were equivalent (*F* test). All statements of statistical significance are based on *P* ≤ 0.05. We used Spearman rank correlations to detect associations between multiple continuous variables.

We handled the problem of self-selection bias by discussing it in every section of this paper. Also, we avoid extrapolating from the data but restrict analyses of percentages and frequencies to within the dataset and note when upward or downward bias might arise from self-selection for extreme cases.

### Ethical note

We conducted no research on hounds or wolves. The data presented are exempt from approvals for research on human subjects because the data were collected and anonymized by another organization (WDNR and SCWC). Institutional review boards and Animal Care and Use Committees are convened to protect human and animal subjects of research not to protect anonymized data. No funding was required for this study and neither author received compensation for the work.

## Results and Discussion

### Wolf-hound interactions (WHI) behaviour, grouping, and phenotype

In 176 case files, we found 140 independent WHI cases during our study period, where we pooled a case reported on the same day and location by different owners into one WHI. Records included 145 killed (83%) and 31 injured (17%) hounds, similar to 71% and 82% reported in Nordic countries (Backeryd 2007); also see (Kojola & Kuittinen 2002). The high percentage of fatalities in both cases might reflect that handlers sometimes took hours to find a distant hound. Therefore, sub-lethal injuries might not be attributed to a WHI if handlers and hounds reunited long after it ended; this effect would bias WHI records upward to more fatalities (see discussion of self-selection bias). Wolf injuries and deaths in WHI were not documented nor reported in case files. Another possible source of missing data might come from misidentifying wolves as the wild interactant, given the availability of compensation for such WHI and not for casualties from other wildlife.

In total, 89% of WHI occurred while hunters reported pursuing black bears, bobcats *Lynx rufus* 6%, coyotes *C. latrans* 4%, raccoons *Procyon lotor* 1%). However, we lack independent data on the animal being pursued by those hounds at the time of WHI. We also lack the relative frequencies of targeting each species with hounds statewide and over time. There was no association between the outcome of WHI categorized as either injury or death and the prey being pursued by hunters (*X*^2^ = 1.9, *p =* 0.75, *df* = 4, *n =* 140). The bear-hound-training period accounted for 62% of WHI, whereas the bear-hunting season accounted for 28% despite being the same length approximately. Outcomes were not associated with month annually (*X*^2^ = 8.5, *p =* 0.38, *df* = 8, *n =* 176, Table 1). These two results suggest that seasonality and wolf reproductive timing did not predict injury or death of hounds, which undermine both the predation and territoriality hypotheses but see further below. We lack information on the frequency with which WHI was initiated by hounds, which could result in different causal hypotheses focused on the motivations of hounds and handlers.

**Table 1.** The number of hounds reported in wolf-hound interactions by breed and month.

| Breed | Jan | Feb | Mar | Apr | May | Jun | Jul | Aug | Sep | Oct | Nov | Dec |
| --- | --- | --- | --- | --- | --- | --- | --- | --- | --- | --- | --- | --- |
| Bluetick | 0 | 1 | 0 | 0 | 0 | 0 | 3 | 6 | 8 | 0 | 0 | 1 |
| Plott | 0 | 0 | 0 | 0 | 0 | 0 | 11 | 20 | 9 | 3 | 0 | 0 |
| Redbone | 0 | 1 | 0 | 0 | 0 | 0 | 4 | 4 | 1 | 0 | 0 | 1 |
| Walker | 0 | 0 | 0 | 1 | 0 | 0 | 16 | 19 | 16 | 3 | 1 | 5 |
| Other | 1 | 2 | 0 | 0 | 0 | 0 | 12 | 15 | 8 | 0 | 0 | 4 |
| **Totals** | **1** | **4** | **0** | **1** | **0** | **0** | **46** | **64** | **42** | **6** | **1** | **11** |

Neither sex nor age of the hounds was associated with the outcome of WHI (sex *X*^2^ = 1.32, *p =* 0.25, *df* = 1, *n =* 151; age *t* = -0.71, *p =* 0.49; variances were equivalent *F* = 0.49). The Treeing Walker Coonhound was the most common breed in WHI (33.3%, *n =* 51), followed by the Plott (27.5%, *n =* 42). There was a suggestive association between breed and outcome (*X*^2^ = 10.7, *p =* 0.03, *df* = 4, *n =* 176). Notably, the Plott fatality frequency of 95.4% was higher than the average 81.2% (Table 1). Plott hounds are the smallest and most vocal hound breed commonly used in Wisconsin for mammals. Small body size has been implicated in the risks and fatalities associated with WHI (Fritts & Paul 1989; Peterson 1995; Backeryd 2007; Kaartinen et al. 2009). The Plott breed is also known for its baying vocalizations, which might alert wolves from a long distance as in (Backeryd 2007; Kaartinen et al. 2009). Small size may make a hound more vulnerable to head and neck bites. Bites to the neck were associated with higher fatality rates in a Scandinavian study (Backeryd 2007). Again, self-selection bias may inflate the involvement of Plott hounds if fatalities were more common for this breed.

Our analysis on hound body site bitten was limited to 109 WHI. We cannot be certain that wolves inflicted every bite because of the above-mentioned delays in discovering hounds and the potential bias created by compensation only for attacks by wolves. Of the 109 carcasses with bite information, 50 provided one bite location (46%), 37 provided two locations (34%), and 22 provided 3 or more locations (20%). Considering all bite locations (*n* = 193), the single most frequent bite site was the neck (33%), followed by back (17%), upper thigh (12%), and chest (10%). We considered bites to the head, shoulders, neck (as opposed to throat), back, and upper thighs as indicative the hound had been lower than its attacker. Those upper body parts were represented in 72% of the 193 bites whereas under-parts (throat, groin, sternum, ribs, lower legs, abdomen) were represented in 28% of bite locations. We found no relationship between body site bitten and outcome, when we separated neck and head bites from others (*X*^2^ = 1.5, *p =* 0.22, *df* = 1, *n =* 66).

Another factor affecting the vulnerability of canids to attack by other canids is numerical superiority. Aggression between wolves and coyotes in Yellowstone National Park had fatal consequences when wolves outnumbered the smaller coyotes, but not when coyotes outnumbered wolves, suggesting that group size exerted less influence than individual body size differences in determining outcomes between canids (Ballard et al. 2003; Merkle et al. 2009). An average of 1.3 hounds were injured or killed per WHI (maximum 5 in a single WHI). The size of the hound group averaged 3.8 *SD* 1.4, *n* = 57. Only 3 (5%) of those WHI involved a single hound in the handler’s control. The number involved in the WHI (2.6 *SD* 1.3, *n* = 47 with 9 (19%) of those WHI involving only 1 hound, were both similar to the number of wolves observed by handlers (2.9 *SD* 1.2, *n* = 15) and similar to the census pack size for the pack blamed by the state or federal agent tasked with verifying the report (2.4 *SD* 1.0, *n* = 19; *n* = 4 included information for both observed and censused). The outcomes were not associated with the number of hounds, number of wolves, or difference between the two in a given WHI by any of the measures of group size or pack size above (Welch test assumes unequal variance, *F* < 0.72, p > 0.41 in every test). Wolves injured or killed hounds in groups with superior numbers in 44% of WHI with such data (*n* = 16). The lack of relationship between pack or group size and outcomes does not support either hypothesis clearly.

Of 80 deaths with data on consumption of a hound carcass, 49% were partially consumed. Of those 80 hounds consumed, 71% occurred July–August and 27% in September– October.. The timing of WHI presents equivocal evidence for both hypotheses. Higher frequencies of WHI occurred during the hound training period in July and August than during the autumn black bear hunt in September and October (Table 1). Elevated risk in July and August might have been associated with the practice of baiting, as wolves visit bear bait sites in search of food (Bump et al. 2013). In Wisconsin, bear bait sites could be legally established as early as April and could last the entire wolf pup-rearing season. Bump et al. (2013) documented that the risk of WHI was three to seven times greater in Wisconsin than in adjacent Michigan, citing the extended bear-baiting period as a probable cause for the much higher rate. That might support the predation hypothesis if bait was more available (or predictable) than wild foods. However, bear baiting was confounded with wolf pup defense predicted by the territoriality hypothesis. The hound training period coincided during the study with wolf use of rendezvous sites or den sites. The consumption of hound carcasses might corroborate the predation hypothesis, but that is not persuasive because consumption was recorded in only approximately half of the WHI and we do not know if the wolves that attacked were the consumers. Nor can we rule out that consumption followed after the primary motivation for aggression in either the hounds or the wolves. The hound carcasses and bite locations provided limited insight. Bites to head, neck, and throat represented 41% of bite locations on hound carcasses. The predation hypothesis might find support from this result because cranio-cervical killing bites are associated with predation by many mammals (Steklis & King 1978; Van Valkenburgh & Ruff. 1987). But the greater height of wolves off the ground then off most hounds would predict such bite locations in any case. In sum, we find equivocal support for both hypotheses. This could imply both are correct or we are missing information, such as whether the hounds were pursuing Wolves. We also do not have information on the body condition of any of the participants, which seems essential data to test the predation hypothesis.

### Handler behaviour and husbandry

Husbandry, such as avoidance of rendezvous sites and use of bells on collars are difficult to evaluate because of a lack of data on prevalence and use. Overall, 55 (31%) records reported whether hounds in WHI wore bells on their collars, with 20 (36%) wearing them and 35 did not (64%). But we have no data on the use of bells among hounds that did not enter the WHI database and the majority of records did not contain any information on devices. Outcomes were not associated with hunter self-reports of affixing bells to collars. Another step handlers might take to protect hounds and wolves would be to release hounds only in low-risk areas. The state did not systematically collect data on whether their warnings about high-risk WCAs were heeded. The WHI records did not contain such information.

### Survey of by-standers

In all, 105 respondents reported adverse incidents with hunting hounds from 51 Wisconsin counties, 4 Michigan counties, and 5 counties from other states. The average number of incidents per county was 2. Seven respondents declined to specify location, but timing distinguished the reports from others’ reports. The 105 respondents reported 119 separate incidents (Table 2). Of the 105, 42% reported the hounds observed were not accompanied by a handler; similarly, 41% reported finding abandoned or lost hounds on their property. In those cases, some respondents reported contacting local animal shelters, law enforcement, or handlers via phone numbers on collars.

**Table 2.** Bystander reports (n=105) of 119 adverse interactions with hounds or their handlers across counties of Wisconsin, USA. Because a respondent might report more than one interaction, we present the counties from most reports 6) to fewest (1), the names of the 51 counties mentioned in reports, the sum of interactions per row, and the maximum number of hounds reported in a single interaction. When multiple reports were filed about the same interaction and the number of hounds differed, we counted only one interaction and averaged the number of hounds, rounding to the higher integer.

| Reports per county | Sum of the interactions | Maximum number of hounds in a single interaction | Counties with interactions reported |
| --- | --- | --- | --- |
| 6 | 15 | 8 | Bayfield, Iron, Sawyer |
| 5 | 5 | 5 | Forest |
| 4 | 4 | 6 | Langlade |
| 3 | 13 | 6 | Chippewa, Dane, Marathon, Polk, Washburn |
| 2 | 18 | 6 | Dodge, Douglas, Dunn, Florence, Lincoln, Oconto, Price, Shawano |
| 1 | 62 | 8 | Ashland, Barron, Brown, Burnett, Calumet,<br>Cheboygan MI, Columbia, Cuyahoga OH, Door,<br>Eau Claire, Fond du Lac, Gogebic, MI, Green,<br>Houghton, MI, Jackson, Kenosha, Kewaunee,<br>Macon, GA, Manitowoc, Marin, CA, Marinette,<br>Milwaukee, Nash, NC, Oneida, Ontonagon, MI,<br>Outagamie, Ozaukee, Rock, Rusk, St. Croix,<br>Sheboygan, Taylor, Trempealeau, Vernon, Vilas,<br>Walworth, Washington, Waukesha, Winnebago,<br>Wood |

Overall, 63% (n=105) described incidents of trespass including hounds running on their property without permission, handlers found on property without permission, or running hounds on property after being denied permission. Of the 105, 18% of respondents described damage to property caused by hounds, including downed fencing, damaged landscaping and gardens, injury to self and livestock, dead wildlife left on property, vandalism, or litter. Of the 105, 11% reported injury to pets or livestock by hounds; 24% reported knowledge of hounds attacking others ’pets or livestock. Of the 105, 8% describe direct encounters with hounds resulting in personal injury or being chased.

Regarding bystander-handler interactions, 31% reported threatening altercations with handlers, including being unwillingly detained by handlers ’trucks on public roads, or their own private driveways. Of the 105, 51% of respondents reported they “feel intimidated by hound handlers,” and 44% feared retaliation from handlers for reporting confrontations to law enforcement.

Finally, approximately a third of respondents described distrust of the information provided by law enforcement officers and also reported filing official complaints upon which no discernible action was taken. Of the 105, 36% believed a conflict of interest existed for law enforcement officers, including game wardens, because of relationships between officers and handlers, or because the officers were believed to hunt with hounds themselves.

### Comparing numbers of hounds from WHI and survey data

Survey respondents reported 2–8 hounds per interaction (average 3.7, mode 2). That average is identical to the average number of hounds that handlers reported in their pack in WHI above despite the different study periods and presumably locations. This seems to be corroborating evidence of accuracy in reports of hound pack sizes in both datasets, as neither set of complainants was aware of the other. Given the rarity of single hounds (5%) in WHI, the bystander reports of >1 hound seem unsurprising. Similarly, bystanders reported >6 hounds in 3 events (8% of reports that include these data) but handlers never reported >6 in their pack after a WHI. The legal limit per handler was ≤6 hounds per handler released from direct control.

## Conclusions

The behaviour of hunters and hounds exploiting mammals has rarely been investigated in North America. Here we fill the gap partly with self-reported data on the behaviour of hounds and wolves associated with aggressive wolf-hound interactions (WHI). We report a shortage of data on wolf injuries. Also, we present self-reported data on the experiences of bystanders exposed to hound behaviour, handler husbandry, and bystander-handler interactions. We report bystanders perceptions or allegations of unlawful actions by handlers or their hounds. Although hound-handlers’ attitudes have been repeatedly and extensively measured, this is the first data on bystanders. We note an absence of systematic observations of behaviour of all participants.

Our inability to find strong support for either the predation hypothesis or the territoriality hypothesis for WHI, which focus on wolf motivation for WHI, suggests either a multi-causal phenomenon or we lack hypotheses for the motivation of hounds or handlers to initiate WHI. Treeing Walker coonhound was the breed most commonly involved in WHI. Most WHI were fatal for hounds. Plott hounds were disproportionally represented among fatalities. We could not confirm or reject a protective role of special collars with bells or defensive features. Nevertheless, we recommend the state obligate veterinary clinics that treat hounds for wildlife injuries to report each such incident so the welfare of hounds and preventive actions taken by handlers can be evaluated by professional veterinarians and wildlife scientists. In light of recent findings that studded leather collars protect cattle against leopards (Khorozyan et al. 2020), recommending more precautions by handlers seems prudent. A majority of WHI involved hounds pursuing black bears compared to other prey but none could confirm what animal the hounds were pursuing when WHI began. Outcomes of WHI (injury or death of hounds) were not associated with the number of wolves observed or censused near the site, or the numerical differences between wolves and hounds, hound age, hound sex, the species of prey targeted by hunters, or the month in which WHI occurred. Regarding numerical superiority, perhaps both the large group sizes of hounds (up to 6 per owner) in Wisconsin and uncertainty about the number of wolves involved, has obscured associations with the fatality rates during WHI.

The survey data and WHI self-reports by handlers are self-selected samples, which cannot be verify directly or used to extrapolate rates, frequencies, or representativeness in space, time, or demography. Nevertheless, the data are the first of their kind to our knowledge and deserve consideration. If even one allegation of unlawful behaviour by handlers was accurate, we recommend attention by law enforcement and hunt managers. We found little financial or non-financial incentive for the self-reports other than bystander dissatisfaction with law enforcement response or value-based disapproval of hounding. By contrast, our other self-selected data set (handler complaints of hound losses in WHI) were motivated by a compensation program that paid for injured or dead hounds. That program also seems to need reform given that no information on harm to wolves was collected despite legal protections for wolves. It might be impossible to verify that the handlers or hounds were acting lawfully at the time of the WHI. Furthermore, the size of packs of hounds (up to 6) and the possibility that multiple handlers can run multiple packs in the same area simultaneously creates a clear danger for wolves and other non-target animals. These results highlight a need for improved regulation, greater oversight and more energetic enforcement of activities involving the use of hounds during hunting.

Our results are also consistent with earlier findings of wildlife crimes (SM 1). Wolf-poaching is the major cause of mortality in US wolf populations (Treves et al. 2017a). Most recently, data reveal that cryptic poaching of Wisconsin wolves rose during hound training, bear-hunting seasons, and deer-hunting seasons (Santiago-Ávila & Treves 2022). The 2021 Wisconsin wolf-hunt saw the most rapid season closure and over-kill in Wisconsin wolf management history with 218 wolves killed in <72 hours with >80% being killed by hunters using hounds; these incidents included at least some hound-inflicted injuries to wolves (Treves et al. 2021).

The topic of comparing self-reports by handlers and self-reports by bystanders raises an issue of expertise, power asymmetries, and research bias. While we have attempted to describe the biases inherent to self-selected respondents in both of our datasets, dismissing these data because they are self reports cannot be justified scientifically either. The larger literature is not so transparent about biases in reporting human-animal interactions. For one, it is routine to publish papers describing adverse effects of wildlife on people but not the benefits. We follow this pattern here. Moreover, Work describing the adverse effects of hunters on wildlife or people is often restricted to so-called bushmeat or distant trade in wildlife parts. Less often, the North American model of hunter behaviour is applauded by various commentators without acknowledging its darker side revealed by data such as ours. Moreover, the human-human conflicts we described from self-selected eye-witnesses exposes another problem in research bias. A common assumption in peer-reviewed commentary on hunting is that hunters have lay knowledge gained through local experience and expertise in their practices (Sandstrom et al. 2009; von Essen et al. 2015). This may favor the views and preferences of hunters over others. But that assumption is unjustified when two local, lay types of expertise are pitted against each other as in some of our study undoubtedly, when hound handlers and the bystanders complaining about them were equally local and both held lay expertise. The power asymmetry reflected in complaints by bystanders described here suggest we need more attention to all participants affected by hunting.

In USA wildlife law, the Endangered Species Act (ESA) and federal court cases surrounding it make clear that some hound handlers are vulnerable to prosecution. First, any “take” of wolves (including harassment, pursuit, injury, killing, etc.) is prohibited under the ESA regardless of whether the perpetrator knew the wild animal harmed was listed (Newcomer et al. 2011). The absence of data revealed by this study indicates that hounding is not adequately regulated. Because wolves were often a federal- or state-listed species during our study, the practice of hound-hunting in wolf pack territories should be prohibited when wolves are a listed species unless it can be regulated to reduce WHI near zero. Prohibitions on non-selective killing methods in the range of endangered species and prohibitions on hunting non-listed species of similar appearance such as coyotes (Newsome et al. 2015), might resolve some WHI. Moreover, if alleged infractions by handlers fall within the statute of limitations, law enforcement should pay heed. Moreover, systematic and repeated failure to investigate or prosecute complaints of illegal activity can make a department vulnerable to lawsuits and the imposition of oversight by higher authorities (e.g., federal consent decrees). Legal jeopardy arises for the agency because the doctrine of prosecutorial discretion may not protect a law enforcement agency from charges of systematic neglect of unlawful activities (Newcomer et al. 2011; Wild Earth Guardians v USA2015).

## Competing Interests

AT and LM perceive no potential competing interests. All funding awarded to AT as of 13 June 2023 are listed here http://faculty.nelson.wisc.edu/treves/archive_BAS/funding.pdf, which we provide for transparency and a CV for disclosure of potential competing interests: http://faculty.nelson.wisc.edu/treves/archive_BAS/Treves_vita_latest.pdf. LM is a retired Associate Professor from the Department of Literature, Leeward College, University of Hawaii and currently a member of: Endangered Species Coalition, Society for the Preservation of Poultry Antiquities, National Wolfwatchers Coalition, Wisconsin Wolf Trackers, Wolf Patrol, Executive Committee, Fox Valley Group, Sierra Club.

## Acknowledgments

No funding was sought or required for this work.

## Supplementary Material 1

Biodiversity may suffer after domestic animals are injured or killed, because their owners may react in several ways detrimental to nature protection efforts. Owners may escalate and kill one or more wild animals, following the incident or for years afterwards. Furthermore, resentments engendered by dangerous wildlife encounters can spread to associates of the involved humans and become broad-based attitudes of intolerance or even preemptive lethal actions against the wildlife. For instance, consider the history of social scientific work done by various authors measuring attitudes to wolves in Wisconsin (Naughton-Treves et al. 2003; Treves et al. 2009; Treves & Martin 2011; Treves et al. 2013; Browne-Nuñez et al. 2015; Hogberg et al. 2015). The first survey in 2001 included complainants who believed they had experienced a wolf attack on their domestic animals, whereas the second survey in 2004 included many more individuals who had not experienced such losses, yet both groups showed decreases in tolerance for wolves when they were resampled in 2009. The interest group least tolerant of wolves was bear hunters who used hounds and the group whose tolerance for wolves declined most over time were men in wolf range who had hunting experience, not those with personal experience of wolf attack on domestic animals (Naughton-Treves et al. 2003; Treves et al. 2009; Treves & Martin 2011; Treves et al. 2013; Hogberg et al. 2015). The prior results on tolerance were paralleled by inclinations to kill wolves illegally (Naughton-Treves et al. 2003; Treves et al. 2013; Browne-Nuñez et al. 2015). Also, attitudes to wolves and inclination to kill wolves illegally were unrelated to the hound handler’s own experience with wolves or their experience with policy interventions relating to WHI such as compensation for hound injuries (Naughton-Treves et al. 2003; Treves et al. 2013). Handlers reported concerns for safety of the hounds and also concerns with access to land and their ability to pursue this pastime in the face of public and political opposition (Browne-Nuñez et al. 2015). Recent research reports that poaching of wolves peaked during seasons of hunting bears and deer and seasons of training hounds (Santiago-Ávila & Treves 2022). Much of the poaching involves concealment or destruction of evidence, which reflects intent to break the law, has repeatedly risen in incidence along with policies that permit some legal wolf-killing in several US wolf populations (SM 1).

These findings indicate that would-be poachers profit from governmental laxity to act unlawfully or that would-be poachers use the cover of legal hunting to act unlawfully (Chapron & Treves 2016a, b; Santiago-Ávila et al. 2020; Louchouarn et al. 2021; Santiago-Ávila & Treves 2022).

## Supplementary Material 2: Survey instrument

### SM 2.0 Adverse Hound Hunting Related Incidents

The Protecting Native Forests and Wildlife Subcommittee of the Sierra Club-Wisconsin Chapter is conducting a survey to determine the frequency and severity of hounding-related experiences you may have encountered where you reside. The survey is anonymous: no identifying information is being requested. A comments section is provided at the end of this form. Completed surveys may be returned to Elizabeth Huntley before conference close, or mailed to: (Redacted for privacy]

What county do you reside in?

2.1 Frequency of Events and observations made
  1. Have you ever observed hunting hounds on your property? YES / NO
  2. Do you recall how many hounds you saw?
    ___1-4
    ___5-8
    ___More than 8
  3. How often have you observed hunting hounds on your property? Once/2-5 times/More than 5 occurrences/Too many times to count
    If YES, were you also able to observe what animal(s) the hounds were pursuing? YES/NO
    List species if YES________________________________________________________
  4. If you have observed hunting hounds on your property, were they accompanied by (a) handler(s)? YES / NO
  5. Have hounds on your property attacked your pets and/or livestock? YES / NO
  6. Are you personally aware of incidents of hounds attacking others’ pets and/or livestock? YES / NO. If “YES”, please describe in comment space below.
  7. Have you ever found abandoned/lost hound(s) in or around your property? YES / NO.
  8. If “YES”, what action did you take if any? Contacted law enforcement/Contacted game warden/ Contacted local humane society or animal control/Took animal in to ___________________ /Other
  Trespassing on private land
  9. Were you ever asked by a hound handler for permission to be on your property for the purposes of hunting? YES / NO
  10. If you were asked, and you replied “No," did the hound handler trespass on your property anyway? YES / NO
  11. If YES, what action did you take? Nothing-fear retribution/Contacted local law enforcement/Contacted local Warden/Contacted WI DNR/Contacted WI Humane Society/Other
  Threats to self, property, pets, livestock by hunting hound handlers
  12. Have you ever experienced damage to your property that you know or suspected was due to hounding activities? YES / NO If YES, please describe in comments below.
  13. Have you ever experienced injury to (a) pet(s) or livestock that you know or suspected was due to hounding activities? YES / NO. If “YES”, please describe (text box provided for explanation).
  14. Have you ever been chased and/or injured by (a) hound dog(s)? YES / NO. If “YES”, please describe with approximate date in comments below.
  15. Have you ever been directly confronted by (a) hound handler(s) and threatened with bodily harm? YES / NO If YES, please describe in comments below.
  16. Have you ever been left an anonymous note that you suspected to be left by hound handlers that threatened you and/or your pets/livestock with harm? YES / NO
  17. Do you personally know of anyone who has experienced real or perceived threats/persecution by hound handlers? YES/NO If YES please describe in comments below.
  18. Do you feel intimidated by hound handlers? YES / NO
  19. Do you fear retribution from hound handlers if you report incidents involving their activity to local law enforcement? YES / NO
2.2 Law enforcement / Game Warden response to complaints, and conflicts of interest
  20. Have you ever contacted local law enforcement, the local game warden, and/or the WDNR about any problem(s) you have experienced with hound handlers? YES / NO
  21. If "YES," what kind of response did you receive?
    ___Warden or Officer responded, but nothing was done.
    ___The Warden/Officer followed up by contacting the suspected hound handler(s), and followed up with you.
    ___The Warden/Officer never responded.
    ___Other.
  22. Do you ever see any local game wardens in the vicinity of your property? Never/On-occasion/All the time
    If YES, is there any particular time of the year when you see them most often? Circle all that are applicable: Deer season/Bear season/Wolf season/Turkey season/Bobcat sea-son/Other
  23. Are you aware of any law enforcement officer and/or game warden who also participates in hounding activities? YES / NO
  24. If "YES," have these been the same individuals who have responded to your complaint(s) about hounding? YES / NO
  25. Are you aware of law enforcement officers / game wardens who have personal relationships with hound handlers? YES/NO If "YES," do you suspect a possible conflict of interest in performing their duties? YES / NO
2.3 Thank you for taking the time to respond to this survey. Please add any additional comments below. You may share your contact information if you wish. The Sierra Club, Wisconsin Chapter will not share your contact information without your permission.

## Supplementary Material 3: Qualitative data received from respondents reporting incidents, anonymized, and edited for clarity

*Hounding should not be allowed all over the state. It’s a free-for-all in the national forests, and it’s wrong. Oconto, WI*

*Please end this cruel sport. Bayfield, WI*
*I do not support running animals with hounds as a hunter. Florence, WI*
*Living in the North woods at this time of year is uncomfortable. Hounders are everywhere, running dogs on bear, blocking roads. Washburn, WI*
*Bear hounds have a negative impact upon wildlife and forest users. I believe hunting bears and wolves with hounds is one of the cruelest “sports” imaginable and separates bear families. Macon, GA*
*I live in the UP and am sick of bear hunters ’lack of respect for land owners and care for their dogs. This is definitely a sport that should be stopped. Houghton, MI*
*I respect the Native cultural relationship with the wolf. We have had a gutted bear (left) on our property. Douglas, WI*
*I have found at least 5 lost hounds along the highway. One had foot injuries. Ontonagon, MI*
*Huge conflict of interest, but we are afraid. Dane, WI*
*I believe about ten years ago, Rep. Frank Boyle attempted to pass an ordinance to protect property owners from uncontrolled hounds. Ashland, WI*
*Bear hunting with hounds promotes unsafe situations for residents in bear country. Burnett, WI*
*Why run hounds in July when wolves have pups and in areas known to be inhabited by packs? Tax payers should not be paying for this. It is a scam. Bayfield, WI*
*Hound hunting is cruel to the hounds and animals being hunted. Gogebic, MI*
*Bear hunters with hounds seem to feel they can trespass. They have a strong enough lobby to pre-vent any repercussions. Chippewa, WI*
*Men standing with trucks and ATVs and radios blocked me on a road on Federal public land. Three cubs ran in front of me. Location withheld*
*People have a right to protect their pets and livestock and not be afraid in their own homes. Fond du Lac, WI*
*Coyote hunters argued with me when we denied permission to hunt on our land. We do not sup-port hounding at all. Manitowoc, WI*
*I’m dismayed this form of cruelty is considered a legal form of “hunting” in Wisconsin. It’s dread-ful and causes harm to a variety of wildlife species, not just the “target.” St Croix, WI*
*I do not approve of hound hunting and believe it should not be practiced. Grant, WI*
*I don’t believe in hound hunting for any animal. Price, WI*
*No animal should be taught to attack another animal and kill it. Location withheld*
*My experience during my 28 years as a municipal clerk is that hound hunting does cause problems. Eau Claire, WI*
*Hound hunting should be stopped. Under no circumstances should those dogs be paid for by tax payers after fights with wolves. Hunters are well aware what will happen to these dogs. Mara-thon, WI*
*The few times I’ve called law enforcement for trespassing, they tell me there’s not much they can do. They’ve told me the hounders have a right to park in front of my driveway and run their dogs on my property. Polk, WI*
*I am very distressed that my state allows hunting of wolves and that the legislature circumvented the necessary period for monitoring. Hunting wolves and bears with hounds is despicable. Dane, WI*
*I hate hunting of any kind with dogs. I don’t understand why it is ok for bear dogs to “train” in our national forests in the summer, harassing wildlife. Price, WI*
*Please, please, let’s do something to stop this! Langlade, WI*
*Hounders ’lack of respect for private property, and just the peace and quiet of my area are being totally disrupted when they run a bear or wolf through the area. Burnett, WI*
*There’s no reason for hound hunting period. Even when they know (wolves have killed) a dog in the area, they still let them go and the hunter gets repaid. I won’t go outside if I see them. Oconto, WI*
*I do not agree with letting dogs chase wild animals for “sport.” It’s disgusting. Wood, WI*
*(They have) no regard for trespassing on private land. Milwaukee, WI*
*Thank you for investigating this problem with wildlife management. Cuyahoga, OH*
*Hounding is unethical. Hounders train their dogs to destroy other animals. Hounders trade, sell, or dump their dogs once they stop performing. Bayfield, WI*
*I am a member of the NRA and Sierra Club. I believe hunting is a right, but the use of dogs to harass, maim, kill other animals is murder. Douglas, WI*
*I live next to hunting hounds and they are very noisy. There have been a lot of complaints, because we have no noise ordinance. I live in the country for nature, not barking dogs. Polk, WI*
*This problem is WAY bigger than me bugging [local law enforcement]…We have a very large dangerous problem with armed gangs of armed criminals breaking laws and terrorizing people. Some people find the level of lawlessness downright frightening. Town of draper has elderly residents that are afraid for their safety. Sawyer, WI*
*They threatened to burn a cabin down not far from my place if the property owner called the war-den. They stole the property owner’s trail cam[era] which had proof of them trespassing with their hounds. County withheld, WI*

## Notes

### Competing Interest Statement

AT and LM perceive no potential competing interests. All funding awarded to AT as of 13 April 2022 are listed here http://faculty.nelson.wisc.edu/treves/archive_BAS/funding.pdf, which we provide for transparency and a CV for disclosure of potential competing interests: http://faculty.nelson.wisc.edu/treves/archive_BAS/Treves_vita_latest.pdf.
LM is a retired from the Department of Literature, Leeward College, University of Hawaii and currently a member of:
Wildlife Conservation Education
Nature Photography
Endangered Species Coalition
Society for the Preservation of Poultry Antiquities
National Wolfwatchers Coalition
Wisconsin Wolf Trackers
Wolf Patrol
Executive Committee, Sierra Club, Fox Valley Group
One Planet. One Life.

### Summary of Updates

Title change, shortened, revisions to avoid the appearance of bias

## References

2015. WildEarth Guardians, et al. v U.S. Department of Justice. U.S. District Court Arizona.

2021. Brown, J. et al. v Kemp, J. et al. U.S. Court of Appeals, 7th circuit.

Backeryd J. 2007. Wolf attacks on dogs in Scandinavia 1995-2005. Ecology Institute. Swedish University of Agricultural Sciences, Grimso.

Ballard WB, Carbyn LN, Smith DW. 2003. Wolf interactions with non-prey. Pages 259-271 in Mech L, and Boitani L, editors. Wolves: behavior, ecology, and conservation. University of Chicago Press, Chicago.

Bowers RR. 1953. The free-running dog menace. Virginia Wildlife 14:5–7.

Bump JK, Murawski CM, Kartano LM, Beyer DE, Roell BJ. 2013. Bear-Baiting May Exacerbate Wolf-Hunting Dog Conflict. PLoS ONE 10.1371/journal.pone.0061708.

Ciucci P, Boitani L. 1998. Wolf and dog depredation on livestock in central Italy. Wildlife Society Bulletin 26:504–514.

Fritts SH, Paul WJ. 1989. Interactions of wolves and dogs in Minnesota. Wildlife Society Bulletin 17:121–123.

Gompper ME 2013. Free-ranging dogs and wildlife conservation. Oxford University Press., Oxford.

Grignolio S, Merli E, Bongi P, Ciuti S, Apollonio M. 2011. Effects of hunting with hounds on a non-target species living on the edge of a protected area. Biological Conservation 144:641–649.

Guagnin M, Perri AR, Petraglia MD. 2018. Pre-Neolithic evidence for dog-assisted hunting strategies in Arabia. Journal of Anthropological Archaeology 49:225–236.

Heberlein TA. 2000. The gun, the dog and the thermos: Culture and hunting in Sweden and the United States. Sweden & America Fall 2000:24–29.

Hristienko H, McDonald JEJ. 2007. Going into the 21st century: a perspective on trends and controversies in the management of the American black bear Ursus 18:72-88.

Kaartinen S, Luoto M, Kojola I. 2009. Carnivore-livestock conflicts: determinants of wolf (Canis lupus) depredation on sheep farms in Finland. Biodiversity and Conservation 18:3503–3517.

Khorozyan I, Siavash G, Mobin S, Soofi M, Waltert M. 2020. Studded leather collars are very effective in protecting cattle from leopard (Panthera pardus) attacks. Ecological S;lutions and Evidence 1:e12013.

Kojola I, Kuittinen J. 2002. Wolf Attacks on Dogs in Finland. Wildlife Society Bulletin 30:498–501.

Kreeger T. 2003. The internal wolf: physiology, pathology, and pharmacology. Pages 192-217 in Mech LD BL, editor. Wolves: behavior, ecology, and conservation. University of Chicago Press, Chicago.

Mech LD 1970. The Wolf: The Ecology and Behavior of an Endangered Species. University of Minnesota Press, Minneapolis.

Merkle J, Stahler D, DW S. 2009. Interference competition between gray wolves and coyotes in Yellowstone National Park. Canadian Journal of Zoology 87:56–63.

Naughton-Treves L, Grossberg R, Treves A. 2003. Paying for tolerance: The impact of livestock depredation and compensation payments on rural citizens’ attitudes toward wolves. Conservation Biology 17:1500–1511.

Newcomer E, Palladini M, Jones L. 2011. The Endangered Species Act v. the United States Department of Justice: How the Department of Justice derailed criminal prosecutions under the Endangered Species Act. Animal Law 17:241–271.

Newsome T, Bruskotter JT, Ripple WJ. 2015. When shooting a coyote kills a wolf: Mistaken identity or misguided management? Biodiversity and Conservation 24:3145–3149.

Olson ER, Treves A, Wydeven AP, Ventura S. 2014. Landscape predictors of wolf attacks on bear-hunting dogs in Wisconsin, USA. Wildlife Research 41:584–597.

Peterson R. 1995. Wolves as interspecific competitors in canid ecology. Pages 315-323 in Carbyn L, Fritts S, and Seip D, editors. Ecology and conservation of wolves in a changing world. Canadian Circumpolar Institute., Edmonton, Alberta, Canada.

Plumer L, Talvi Tn, Männil P, Saarma U. 2018. Assessing the roles of wolves and dogs in livestock predation and suggestions for mitigating human-wildlife conflict and conservation of wolves. Conservation Genetics https://doi.org/10.1007/s10592-017-1045-4.

Pulliainen E. 1965. Studies on the wolf (Canis lupus) in Finland. Annales Zoologici Fennici 2:215–259.

Roosevelt TD 1902. Hunting the Grisly and Other Sketches. G. P. Putnam’s Sons, New York.

Ruid DB, Paul WJ, Roell BJ, Wideven AP, Willging RC, Jurewicz RL, Lonsway DH. 2009. Wolf–human conflicts and management in Minnesota, Wisconsin, and Michigan. Pages 279-295 in Wydeven AP, Van Deelen TR, and Heske EJ, editors. Recovery of Gray Wolves in the Great Lakes Region of the United States: An Endangered Species Success Story. Springer, New York.

Ruprecht J, Ausband D, Mitchell M, Garton M, Zager P. 2012. Homesite attendance based on sex, breeding status, and number of helpers in gray wolf packs. Journal of Mammalogy 93:1001–1005.

Sandstrom C, Pellikka J, Ratamaki O, Sande A. 2009. Management of Large Carnivores in Fennoscandia: New Patterns of Regional Participation. Human Dimensions of Wildlife 14:37–50.

Santiago-Ávila FJ, Treves A. 2022. Poaching of protected wolves fluctuated seasonally and with non-wolf hunting. Scientific Reports 12:e1738

SAS Institute I. 2013. JMP 11.0, Cary, North Carolina..

Steklis HD, King GE. 1978. The craniocervical killing bite: Toward an ethology of primate predatory behavior. Journal of Human Evolution 7:567–581.

Stillfried M, Belant J, Svoboda N, Beyer D, Kramer-Schadt S. 2015. When top predators become prey: Black bears alter movement behaviour in response to hunting pressure. Behavioural Processes 120:30–39.

Treves A, Artelle KA, Darimont CT, Parsons DR. 2017a. Mismeasured mortality: correcting estimates of wolf poaching in the United States. Journal of Mammalogy 98:1256–1264.

Treves A, Jurewicz RL, Naughton-Treves L, Rose RA, Willging RC, Wydeven AP. 2002. Wolf depredation on domestic animals: control and compensation in Wisconsin, 1976-2000. Wildlife Society Bulletin 30:231-241.

Treves A, Jurewicz RL, Naughton-Treves L, Wilcove D. 2009. The price of tolerance: Wolf damage payments after recovery. Biodiversity and Conservation 18:4003–4021.

Treves A, Kapp KJ, Macfarland DM. 2010. American black bear nuisance complaints and hunter take. Ursus 21:30–42.

Treves A, Langenberg JA, López-Bao JV, Rabenhorst MF. 2017b. Gray wolf mortality patterns in Wisconsin from 1979 to 2012. Journal of Mammalogy 98:17–32.

Treves A, Santiago-Ávila FJ, Putrevu K. 2021. Quantifying the effects of delisting wolves after the first state began lethal management. PeerJ 9: e11666.

Van Ballenberghe V, Mech L. 1975. Weights, growth, and survival of timber wolf pups in Minnesota. Journal of Mammalogy 56:44–63.

Van Valkenburgh B, Ruff. CB. 1987. Canine tooth strength and killing behaviour in large carnivores. Journal of Zoology 212:379–397.

von Essen E, Hansen HP, Kallstrom HN, Peterson MN, Peterson TR. 2015. The radicalisation of rural resistance: How hunting counterpublics in the Nordic countries contribute to illegal hunting. Journal of Rural Studies 39:199–209.

Willging R, Wydeven AP. 1997. Cooperative wolf depredation management in Wisconsin. Pages 46-51 in Lee CD, and Hygnstrom SE, editors. Thirteenth Great Plains Wildlife Damage Control Workshop Proceedings. Kansas State University Agricultural Experiment Station and Cooperative Extension Service, Lincoln, NE.

Wydeven AP, Treves A, Brost B, Wiedenhoeft JE. 2004. Characteristics of wolf packs in Wisconsin: Identification of traits influencing depredation. Pages 28-50 in Fascione N, Delach A, and Smith ME, editors. People and Predators: From Conflict to Coexistence. Island Press, Washington, D. C.

## References

Browne-Nuñez C, Treves A, Macfarland D, Voyles Z, Turng C. 2015. Tolerance of wolves in Wisconsin: A mixed-methods examination of policy effects on attitudes and behavioral inclinations. Biological Conservation 189:59–71.

Chapron G, Treves A. 2016a. Blood does not buy goodwill: allowing culling increases poaching of a large carnivore. Proceedings of the Royal Society B 283:20152939.

Chapron G, Treves A. 2016b. Correction to ‘Blood does not buy goodwill: allowing culling increases poaching of a large carnivore’. Proceedings of the Royal Society B Volume 283:20162577.

Hogberg J, Treves A, Shaw B, Naughton-Treves L. 2015. Changes in attitudes toward wolves before and after an inaugural public hunting and trapping season: early evidence from Wisconsin’s wolf range. Environmental Conservation 43:45–55.

Louchouarn NX, Santiago-Ávila FJ, Parsons DR, Treves A. 2021. Evaluating how lethal management affects poaching of Mexican wolves (registered report). Royal Society Open Science 8:200330.

Santiago-Ávila FJ, Chappell RJ, Treves A. 2020. Liberalizing the killing of endangered wolves was associated with more disappearances of collared individuals in Wisconsin, USA. Scientific Reports 10:13881.

Treves A, Martin KA. 2011. Hunters as stewards of wolves in Wisconsin and the Northern Rocky Mountains, USA. Society and Natural Resources 24:984–994.

Treves A, Naughton-Treves L, Shelley VS. 2013. Longitudinal analysis of attitudes toward wolves. Conservation Biology 27:315–323.

